# Mutagenicity of folic acid deficiency and supplementation is tissue-specific and results in distinct mutation profiles

**DOI:** 10.1101/2020.07.27.223552

**Authors:** Stephanie Diaz G., Danielle P. LeBlanc, Remi Gagné, Nathalie A. Behan, Alex Wong, Francesco Marchetti, Amanda J. MacFarlane

## Abstract

Cancer incidence varies by tissue due to differences in environmental risk factor exposure, gene variant inheritance, and lifetime number of stem cell divisions in a tissue. Folate deficiency is associated with increased risk for colorectal cancer (CRC) and acute lymphocytic leukemia. Conversely, high folic acid (FA) intake has been associated with higher CRC risk. However, the mutagenic potential of FA intake in different tissues has not been characterized. Here we quantified mutations in folate-susceptible somatic tissues, namely bone marrow and colon, from the same MutaMouse mice and determined the FA-induced mutation profiles of both tissues using next generation sequencing. FA-induced mutagenesis was tissue- and dose-specific: FA deficiency increased mutant frequency (MF) in bone marrow while FA supplementation increased MF in colon. Analyses of mutation profiles suggested that FA interacted with mutagenic mechanisms that are unique to each tissue. These data illuminate potential mechanisms underpinning differences in susceptibility to FA-related cancers.

## Introduction

Mutations in somatic cells accumulate through life and are influenced by endogenous and exogenous processes^1^. Cells that incur mutations that confer a selective advantage demonstrate preferential growth and survival, increasing the risk for cellular transformation and cancer^1,2^. Cancer is common, accounting for ~1 in every 6 deaths worldwide; however cancer incidence varies considerably by tissue^3^. Tissue differences in cancer incidence are attributable to variable exposures to environmental risk factors, inherited genetic variation, and to differences in the lifetime number of cell divisions of a tissue’s stem cells^4^. Environmental risk factors, such as chemical contaminants, self-care products, and nutrients, are associated with tissue-specific cancers, seemingly contributing to carcinogenesis in one tissue while having little or no effect in another, suggesting tissue-specific responses^5–9^

Folate deficiency due to inadequate intake or polymorphisms in genes related to folate metabolism is associated with the risk for specific cancers, such as acute lymphocytic leukemia (ALL) and colorectal cancer (CRC)^10–13^. In the case of CRC, FA supplementation is associated with both reduced and increased risk; differences in the effect of supplementation may depend on dose, duration, cancer stage at intervention, and baseline folate status, among other contributing factors^14–19^. While the association between FA intake and tissue-specific cancer risk is well-established, the mechanism(s) underpinning these relations is not understood.

Folate-mediated one-carbon (1C) metabolism is required for the *de novo* synthesis of purines and thymidine (dTMP), and the synthesis of methionine, a precursor of the universal methyl donor, S-Adenosylmethionine (AdoMet)^20^. Disruption of any of the folate-dependent biosynthetic pathways has implications for mutagenesis and genome stability. Impairment of *de novo* purine synthesis causes nucleotide pool imbalances and impairs DNA repair thereby inducing mutations^21,22^. Insufficient dTMP synthesis results in uracil misincorporation into DNA leading to DNA double strand breaks. Disruption of AdoMet production reduces cellular methylation capacity; altered genomic methylation patterns can cause genome instability and modify gene expression^23^. These processes contribute to cellular transformation and cancer risk.

Here, we hypothesized that FA intake contributes to mutagenicity, a proxy for cancer risk, in a tissue-specific manner. Using the MutaMouse, an established transgenic mouse model used for assessing *in vivo* mutagenicity^24^, we compared the effect of dietary FA deficiency and supplementation on the mutant frequency (MF) in the colonic epithelium and bone marrow. We then characterized the mutational profiles in each tissue using next generation sequencing (NGS). We demonstrate that the mutagenic potential of FA is tissue- and dose-specific with FA deficiency inducing mutations in the bone marrow and supplementation inducing mutations in the colon. Each tissue displayed distinct mutational profiles with similarity to known human cancer mutation profiles and associated with specific mechanisms. Our data identify candidate causal mechanisms underpinning the tissue-specificity of folate-associated cancers.

## Results

### Folic acid supplementation increases mutant frequency in the colon

Beginning at 5 weeks of age, MutaMouse male mice were fed one of three FA diets: 0 mg FA/kg (deficient), 2 mg FA/kg (control) or 8 mg FA/kg (supplemented) and maintained on the diets for a total of 20 weeks (Supplementary Figure 1). These diets represent physiologically relevant intakes of FA. The deficient diet (0 mg FA/kg) represents an inadequate dietary FA intake, resulting in tissue folate depletion, higher circulating homocysteine and higher micronucleous frequency in red blood cells (RBCs)^25,26^. The control diet (2 mg FA/kg) is an adequate dietary FA intake for rodents, as recommended by the American Institute of Nutrition. It corresponds with the Recommended Dietary Allowance (RDA) for adult humans of 400 μg dietary folate equivalents per day^27–29^. The supplemented diet (8 mg FA/kg) approximates a dietary FA intake of 1.6 mg per day in adult humans, more than the Tolerable Upper Intake Level (UL) of FA of 1 mg/day, which is the highest daily intake likely to pose no risk of adverse health effects^28^.

At the 10-week timepoint, mice were treated with saline or *N*-ethyl-*N*-nitrosurea (ENU), a known mutagen^30^, to determine the effect of FA alone and its interaction, if any, with ENU on mutations. Recovery of the *lacZ* reporter transgene and *in vitro* positive selection for mutant copies of the transgene^31^ allowed for the measurement of colon MF (Table 1, Figure 1). Among the saline treated mice, those fed the FA supplemented diet had a 1.5-fold higher MF (MF= 14.0 x 10^-5^) compared to mice fed the FA control diet (MF=9.6 x 10^-5^; p = 0.001) and 1.3-fold higher MF compared to those fed the FA deficient diet (MF=10.6 x 10^-5^; p = 0.008). The MF did not differ between mice fed the FA control and deficient diets. As expected, the MF was higher in mice treated with ENU compared to those treated with saline regardless of diet (ENU effect, p<0.00001) but there were no diet-dependent differences among the ENU-treated mice. Because mutagenesis is a proxy for cancer risk, our results suggest that FA supplementation alone may increase the risk of CRC in some cases. Given the complex aetiology of cancer, the heterogeneous observations from randomized controlled trials (RCT) and the smaller magnitude of effect on mutations compared to a strong mutagen, the impact of FA supplementation on cancer, if any, is likely context specific^14,15,17,19,32,33^.

**Figure 1.**
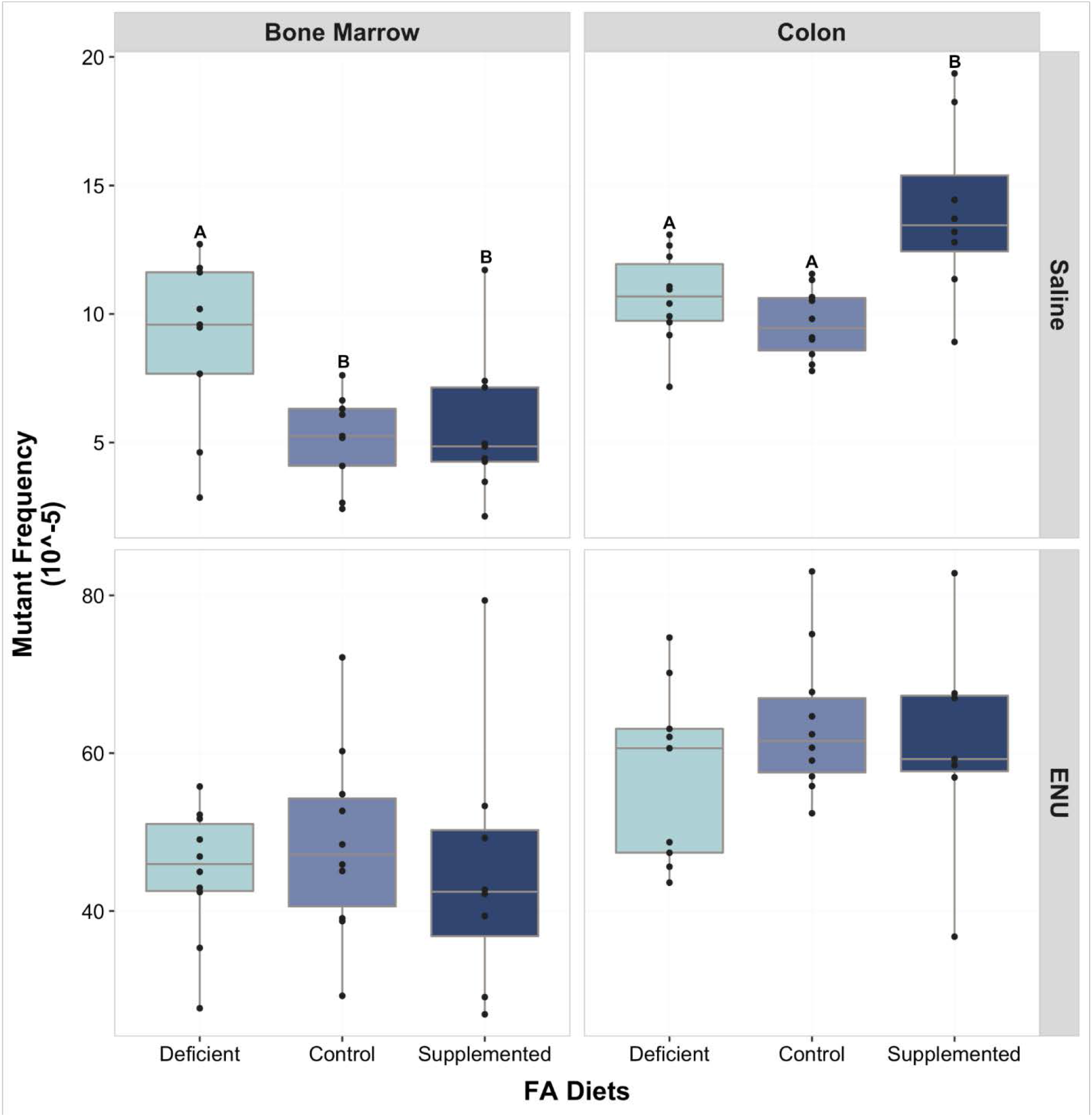
Bone marrow and colon folic acid (FA)-induced *lacZ* MF. *lacZ* mutant frequencies of bone marrow and colon in male mice fed a FA deficient, control or supplemented diet treated with saline and ENU. Note the difference in the Y-axis scale between the saline and ENU groups. ENU effect was significantly higher compared to saline (p = 0.00001). (Two-way ANOVA, Tukey HSD and One-way ANOVA, Holm-Sidak p≤0.05 considered significantly different). Diets: Control = 2 mg FA/kg; Deficient= 0 mg FA/kg; Supplemented = 8 mg FA/kg. Treatment: ENU = N-ethyl-N-nitrosurea (50 mg/kg). n=8-10/diet/treatment.

**Table 1:**
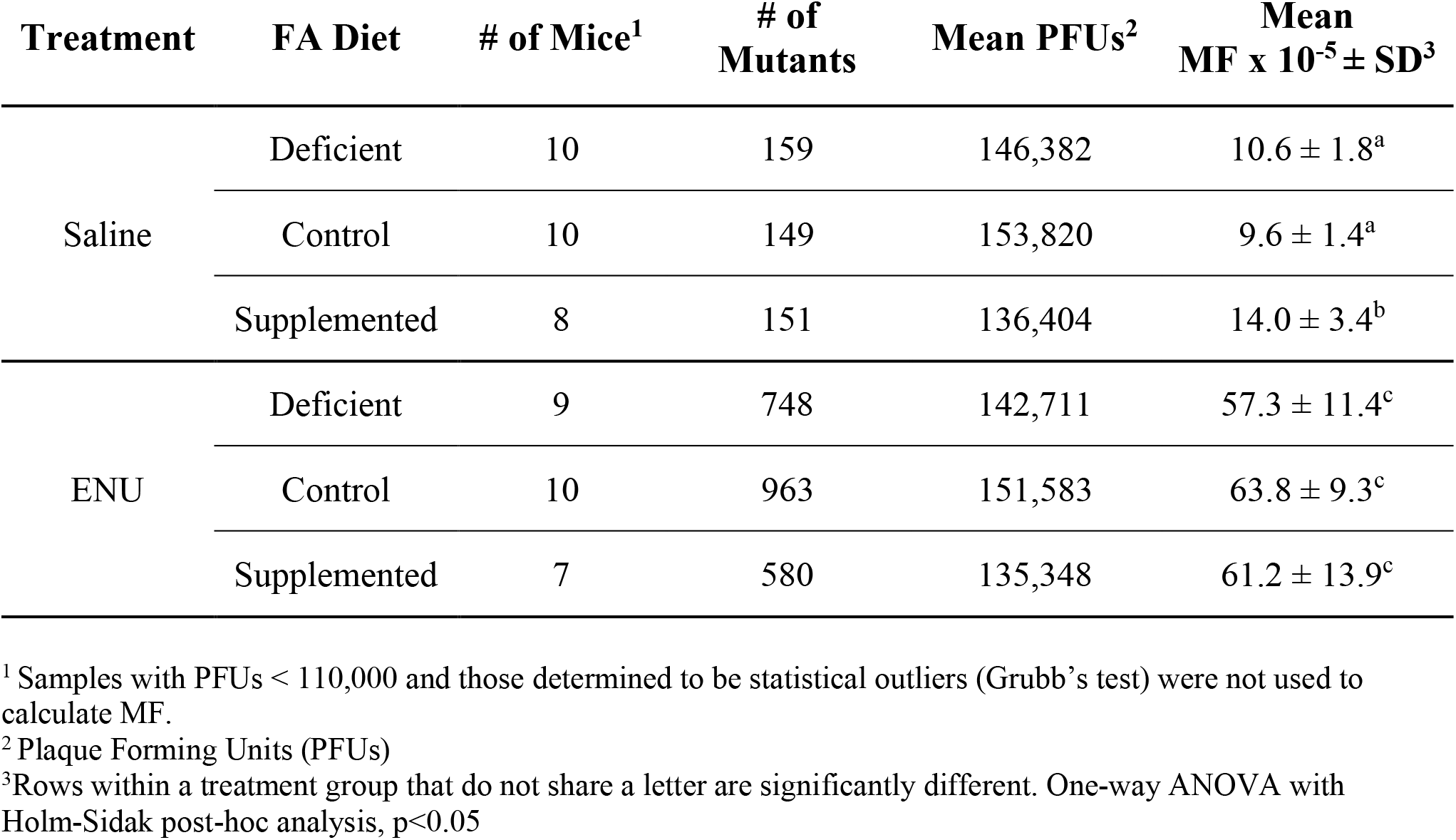
Summary of colon mutant frequencies in mice fed the folic acid (FA) diets and treated with saline or ENU.^1^

| Treatment | FA Diet | # of Mice<sup>1</sup> | # of Mutants | Mean PFUs<sup>2</sup> | Mean MF x 10<sup>-5</sup> ± SD<sup>3</sup> |
| --- | --- | --- | --- | --- | --- |
| Saline | Deficient | 10 | 159 | 146,382 | 10.6 ± 1.8<sup>a</sup> |
|  | Control | 10 | 149 | 153,820 | 9.6 ± 1.4<sup>a</sup> |
|  | Supplemented | 8 | 151 | 136,404 | 14.0 ± 3.4<sup>b</sup> |
| ENU | Deficient | 9 | 748 | 142,711 | 57.3 ± 11.4<sup>c</sup> |
|  | Control | 10 | 963 | 151,583 | 63.8 ± 9.3<sup>c</sup> |
|  | Supplemented | 7 | 580 | 135,348 | 61.2 ± 13.9<sup>c</sup> |

<sup>1</sup> Samples with PFUs < 110,000 and those determined to be statistical outliers (Grubb's test) were not used to calculate MF.
<sup>2</sup> Plaque Forming Units (PFUs)
<sup>3</sup> Rows within a treatment group that do not share a letter are significantly different. One-way ANOVA with Holm-Sidak post-hoc analysis, p<0.05

### Folic acid-induced MF is tissue-specific

We compared the colon MF with previous data collected on the bone marrow from the same saline-treated mice^34^. Bone marrow MF was re-analyzed using the same statistical criteria used for the colon. The MF in colon of mice on the FA control diet was significantly higher than that of bone marrow (p = 0.001; Figure 1), consistent with previously reported historical background MF values for these tissues^31,35^. In contrast to the observed MF in colon, the bone marrow of mice fed the FA deficient diet had a 1.7-fold higher MF (MF= 8.9 x 10^-5^) than that of mice fed the control diet (MF= 5.1 x 10^-5^; p = 0.02), and 1.6-fold higher MF than mice fed the supplemented diet (MF= 5.6 x 10^-5^; p = 0.03). The MF in bone marrow did not differ between mice fed the control and supplemented diets. Given that both tissues were obtained from the same mice, these results indicate that the effect of FA intake on MF is tissue specific.

### FA-induces different mutation profiles in colon and bone marrow

We used NGS to sequence the *lacZ* transgene from mutant plaques (positive selection) to the determine the FA-induced mutation profile in the colon and bone marrow of saline treated mice. Mutant plaques from the bone marrow of 29 mice (463 plaques) and the colon of 28 mice (452 plaques) representing all three diet groups were sequenced in duplicate which resulted in the identification of 400 and 361 recurrent mutations representing 149 and 165 independent mutations in bone marrow and colon, respectively (Supplementary Table 1). The FA-induced mutation profiles for both tissues are shown in Figure 2. Since the control diet represents adequate folate intake, mutation types in the control diets of both tissues were attributed to spontaneous mutations. Spontaneous mutation profiles were dominated by G:C➔A:T transitions, representing ~60% and ~47% of the mutations in the bone marrow and colon of control mice, respectively. The proportion of G:C➔A:T transitions at CpG sites, which can occur spontaneously due to methylcytosine deamination^36^, did not differ among the diets in either tissue. In the colon, while the MF was higher in mice fed the FA supplemented diet, the overall proportions of the observed individual mutation types (G:C➔A:T, A:T➔G:C, G:C➔C:G, G:C➔T:A, A:T➔T:A, A:T➔C:G, deletions and insertions) were similar among the diets. Conversely, in the bone marrow, mice fed the FA deficient diet had a significantly higher proportion of G:C➔C:G transversions compared to those fed the FA control (p=0.016, one-way ANOVA, Tukey HSD) and supplemented (p=0.02, one-way ANOVA, Tukey HSD) diets; the proportions of the other mutation types did not differ by diet.

**Figure 2.**
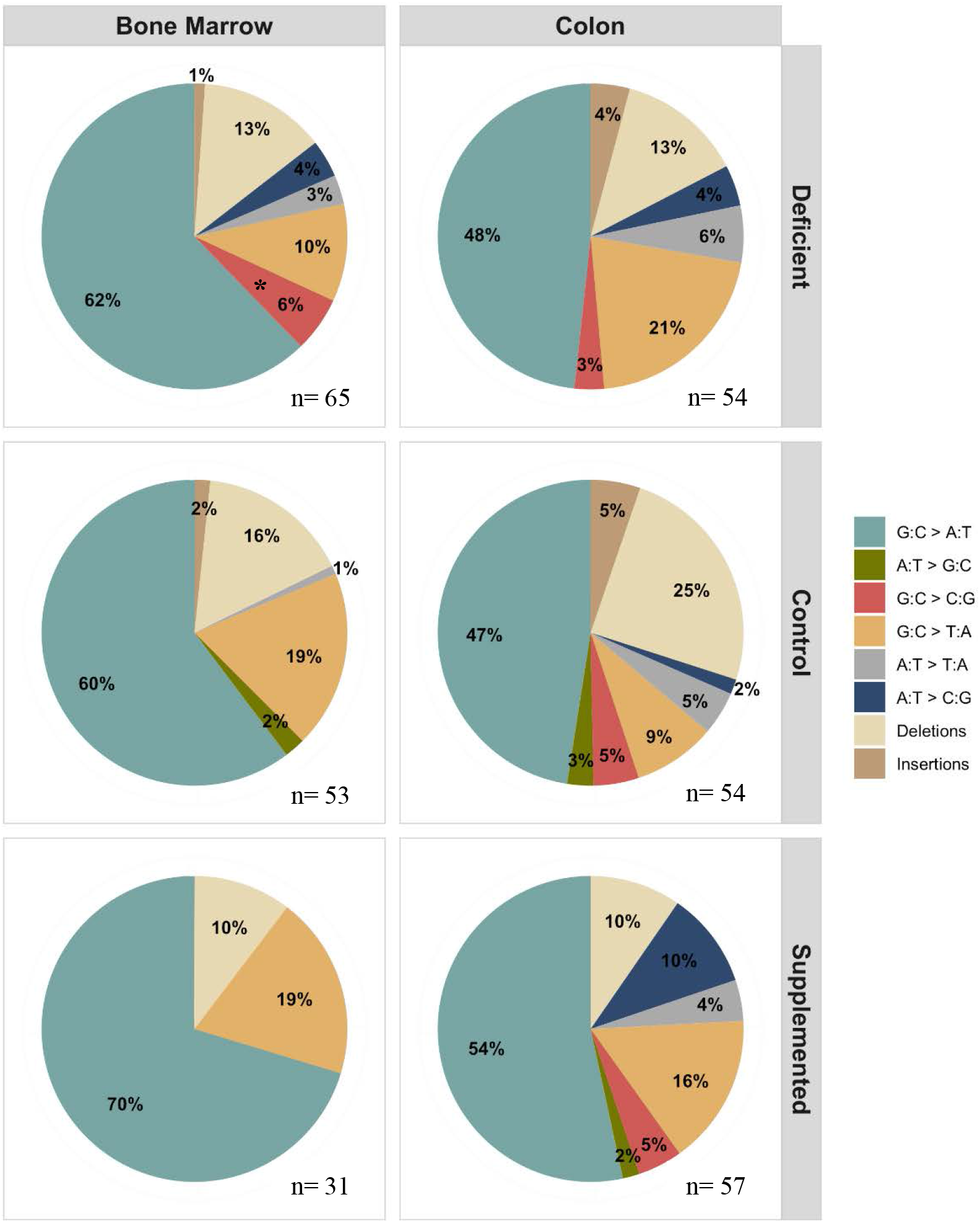
Bone marrow and colon mutation profiles. Proportional representation of *lacZ* mutations in male mice treated with saline and fed one of three folic acid (FA) diets. In the bone marrow, G:C > C:G transversions represent a larger proportion of induced mutations in mice fed the FA deficient diet compared to mice fed the FA control or supplemented diets (One way ANOVA, Tukey HSD, *p<0.05). The relative proportions of the different mutation types did not differ among the diet groups in the colon. Diets: Control = 2 mg FA/kg; Deficient= 0 mg FA/kg; Supplemented = 8 mg FA/kg. n = Total number of unique mutations.

### FA-induced mutational signatures of colon and bone marrow

We next determined the FA-induced mutational signatures in the two tissues (Figure 3). Single base substitution (SBS) signatures account for the sequence context in which the substitution (mutation) occurred. As such, SBS signatures contain 96 mutation types based on the six possible base substitutions (G:C➔A:T, A:T➔G:C, G:C➔C:G, G:C➔T:A, A:T➔T:A, A:T➔C:G) within a trinucleotide sequence where the mutated base is represented by the pyrimidine of the base pair, and includes the bases located immediately 5’ and 3’ to the mutated base^37^. Like the mutation profiles, the bone marrow and colon mutational signatures differed by tissue and reflected the mutation types characterized by each tissue’s mutation profile. For example, while the C>T transition was the most predominant type of mutation observed for all diets in both tissues, the sequence context in which it occurred differed between the tissues. In bone marrow, it occurred most frequently within the T[C>T]G trinucleotide, while in the colon the C>T substitution was more evenly distributed among all 16 possible trinucleotides (Figure 3).

**Figure 3.**
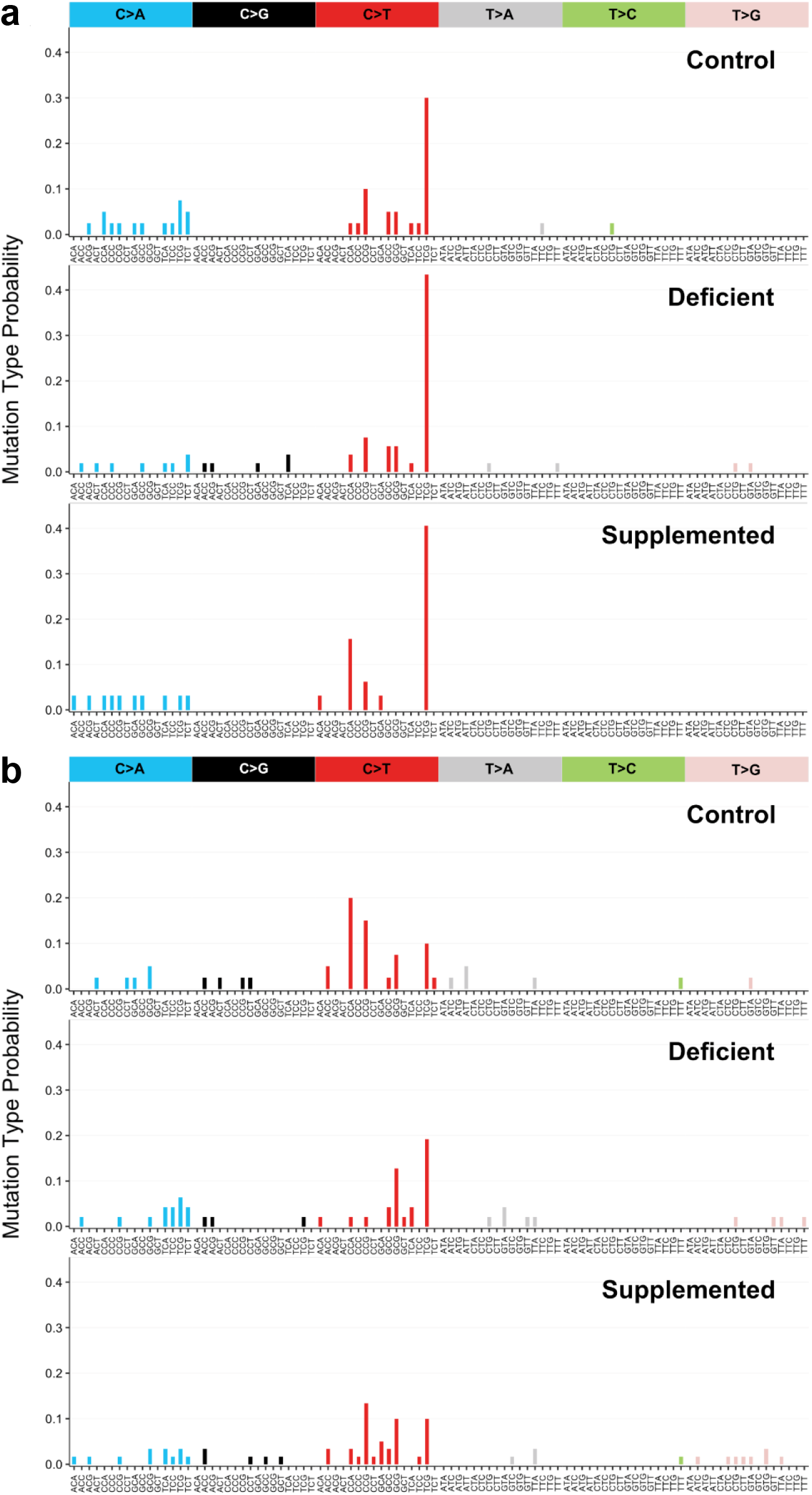
*lacZ* folic acid (FA)-induced mutational signatures of a) bone marrow and b) colon among all FA diets. Bars represent the relative frequency of the mutation in the context of the trinucleotides on the x-axis.

The catalogue of somatic mutations in cancer (COSMIC) contains 49 SBS signatures observed in human cancers, some with known aetiologies^37^. In order to identify candidate mechanisms underpinning the observed tissue- and diet-specific mutations, we identified COSMIC SBS signatures that were most similar to our FA-induced signatures using cosine similarity testing (Figure 4; Supplementary Figure 2). The ratio of trinucleotide frequencies in the *lacZ* transgene to those of the human genome was used to normalize the COSMIC SBS signatures to *lacZ* trinucleotide frequencies^38^. In this exploratory analysis, only COSMIC SBS signatures with a cosine similarity > 0.5 were considered (Figure 4; Supplementary Tables 2 and 3).

**Figure 4.**
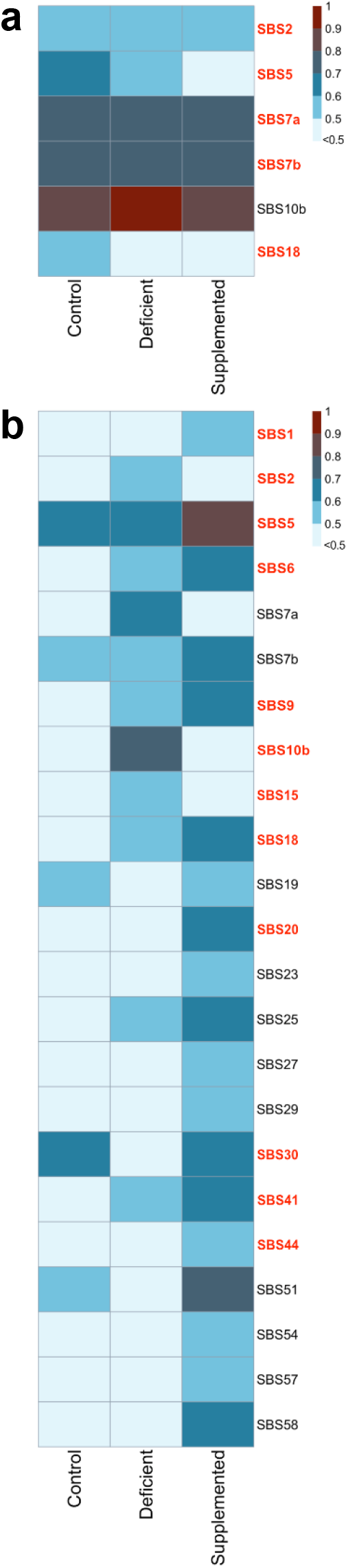
Cosine similarities heatmap of folic acid (FA)-induced mutational signatures and COSMIC SBS signatures (minimum cosine similarity >0.5) in a) bone marrow and b) colon. SBS in red are signatures observed in cancers that develop in each corresponding tissue, namely acute lymphocytic leukemia and colorectal cancer.

The mutation profiles of bone marrow and colon from mice on the FA control diet were associated with different repertoires of SBS signatures. In fact, there were six and five SBS signatures that had cosine similarity values >0.5 with the mutation profile of the bone marrow (Figure 4a; Supplementary Table 2) and colon (Figure 4b; Supplementary Table 3), respectively. Among these SBS signatures, only SBS5 and SBS7b were shared between the two tissues. In bone marrow, no signature was identified that was unique to the FA deficient diet for which we observed a higher MF (Figure 4a, Supplementary Table 2). All FA diet signatures were associated with SBS 2 (cosine ≥0.53), SBS 7a (cosine ≥0.75), SBS 7b (cosine ≥0.74) and SBS 10b (cosine ≥0.89), with the latter having cosine similarity values >0.85 with all three FA diets and reaching the highest value with the FA deficient diet signature (cosine of 0.95) (Figure 4, Supplementary Table 2, Supplementary Figure 3). These four SBS signatures are all characterized by C>T transitions. FA control and deficient signatures were associated with SBS 5 (cosine ≥0.52), while only the FA control diet was associated with SBS 18. Of note, SBS 2, 5 and 7a,b signatures were previously identified in ALL (cancer.sanger.ac.uk)^39^.

In the colon, the FA supplemented diet signature had a cosine similarity values of 0.5-0.8 with 9 SBS signatures (SBS 1, 20, 23, 27, 29, 44, 54, 57 and 58), of which SBS 1, 20, and 44 were previously identified in CRC (cancer.sanger.ac.uk)(Figure 4, Supplementary Figure 4, Supplemental Table 3)^39^. The FA deficient and supplemented diet signatures had cosine similarity ≥0.50 with five SBS signatures (SBS 6, 9, 18, 25, 41); however, the FA supplemented diet signature had higher similarity than the deficient diet for all five signatures (Supplemental Table 3). Four of these five SBS signatures (SBS 6, 9, 18 and 41) had previously been identified in CRC (cancer.sanger.ac.uk)^39^. All FA diet signatures had a cosine similarity ≥0.55 with SBS 5 and 7b (Supplementary Table 3).

These results show that bone marrow and colon have both common and unique SBS signatures that contribute to their spontaneous mutation profiles and that FA interacts with these signatures to result in a tissue- and dose-specific mutagenic response.

## Discussion

In this study, we determined the mutagenic potential of FA intake *in vivo* in two highly proliferative somatic tissues that are susceptible to folate-associated cancers. We demonstrate that the effect of FA intake on MF in bone marrow and colon were tissue- and dose-dependent with opposite effect directions. Moreover, we observe that FA intake interacts with the spontaneous repertoire of SBS signatures of each tissue to result in a tissue- and dose-specific mutagenic response. Our data reveal that FA can have distinct, and in this case even opposing, effects on mutations between tissues within an individual, indicating that the tissue context influences the impact of FA intake on cancer risk.

In bone marrow, we observed a significantly higher MF in mice fed the FA deficient diet (Figure 1), which was not observed in the colon. While both tissues are considered highly proliferative, bone marrow has a higher kinetic turnover and cell renewal rate than colon^40–42^. As such, bone marrow has relatively higher nucleotide synthesis requirements, thereby making it potentially more susceptible to mutations in a folate deficient environment. The bone marrow mutation profile showed a significantly higher proportion of G:C➔C:G transversions in mice fed the FA deficient diet compared to those fed the FA control and supplemented diets, suggesting that the high MF in those fed the deficient diet was driven by G:C➔C:G transversions, a mutation type observed in ALL. The FA deficient diet mutational signature was similar to SBS 2, which is characterized by C>T transitions and has been observed in ALL. SBS 2 is closely associated with SBS 13, which is characterized by C>G transversions^37^, the type of mutation that was significantly higher in the mutation profile of the FA deficient diet. Both SBS 2 and 13 are attributed to the activity of AID/APOBEC enzymes^39^. The AID/APOBEC enzyme family is involved in DNA cytosine deamination (C>U), the repair of which can result in a C>T transition or a C>G transversion^43^. During DNA replication, most polymerases insert an adenine opposite to a uracil in the complementary DNA strand, resulting in a C>T transition following subsequent DNA replication^43^. However, the action of a translesion synthesis DNA polymerase such as REV1, a deoxycytidyl transferase, inserts a cytosine opposite to a uracil, resulting in a C>G transversion^43,44^. Because folate deficiency increases the level of uracil in DNA^45,46^, we propose that it is through the repair of the misincorporated uracil by REV1, present in bone marrow stem cells^47^, that leads to C>G transversions in bone marrow.

In the colon, we observed that mice fed a FA supplemented diet had a significantly higher MF compared to mice fed the control and deficient diets (Figure 1), in contrast to the bone marrow in which the mice fed the FA deficient diet had a significantly higher MF. However, there were no diet-dependent differences in the relative proportions of mutation types in the colon. These results suggest that the mechanism of action underpinning the mutagenic potential of FA supplementation might be the establishment of a mutation permissive context in which the types of mutations that are induced under adequate folate status are all more likely to occur, resulting in an overall increase in the number of mutations but not in any specific type being preferentially induced.

The different effect of FA supplementation in the colon with respect to the bone marrow may be a consequence of direct exposure to FA. Unlike the bone marrow, where cells are exposed primarily to 5-methylTHF, the colonic epithelium would be exposed primarily to FA. FA needs to be sequentially reduced to dihydrofolate (DHF) and then to tetrahydrofolate (THF) by dihydrofolate reductase (DHFR) prior to entering 1C metabolism^48,49^; it is through this pathway that high concentrations of FA in the colon could potentially disrupt 1C metabolism and increase mutations. First, FA could saturate the enzymatic activity of DHFR resulting in the depletion of THF, limiting its availability for 1C metabolism^50^. Second, accumulation of DHF is proposed to act as an inhibitor of key enzymes (i.e.; thymidylate synthase, AICAR transformylase and MTHFR) required for *de novo* dTMP, purine and methionine synthesis^51–53^. Therefore, overall disruption of the folate-dependent biosynthetic pathways by either DHFR saturation or increased DHF concentration could underpin the increase in all types of mutations that was observed in the colon.

Our exploratory analysis to identify COSMIC SBS signatures that are associated with the FA supplemented diet mutational signature in the colon (Supplementary Figure 4) suggests other possible mechanisms. Interestingly, three of the SBS signatures that have been observed in CRC (SBS 6, 20, 44) demonstrated 35-60% higher cosine similarity values with the FA supplemented diet compared to the control diet. These signatures are associated with deficient DNA mismatch repair (MMR). MMR deficiency occurs in ~15% of sporadic CRC cases^54,55^. Disruption of MMR can result from silencing of *MLH1*, an MMR gene, by hypermethylation of its promoter^56,57^ and high serum folate concentration has been positively correlated with *MLH1* promoter methylation^58^. Given that 1C metabolism supports cellular methylation capacity by the production of AdoMet, we propose that FA supplementation could lead to an increased availability of AdoMet in the colon which could promote the hypermethylation of *MLH1*. Additional studies are needed to determine whether FA supplementation is enough to cause hypermethylation of the *MLH1* gene, thereby disrupting MMR, and whether it, in turn, impacts mutagenesis in the colon.

The lack of an effect of the FA deficient diet on mutagenesis in the colon was unexpected given previous observations that suggest an association between folate deficiency and CRC risk. The presence of folate-producing bacteria in the colon could have provided an alternative source of folate for colonocytes, thus mitigating the effect of dietary deficiency on mutations, at least for the duration of this experiment^59,60^. Other factors, such as sex and mouse strain, could also have impacted the effect of diet and this one study does not preclude the possibility that FA deficiency under certain conditions induces mutations in the colon.

This study was strengthened by the use of an established transgenic mouse model for *in vivo* mutagenicity^24^ and the implementations of an approach that allowed for quantification mutations and their molecular characterization by sequencing^61^. This study also had a number of limitations including the limited number of FA doses, the duration of the experiment, and the use of a single sex and mouse strain. Although we did not measure folate vitamers in circulation or in tissues, the plasma folate was measured indicating that the folate status of the mice reflected the diet FA content^34^. This study could have been strengthened by sequencing a larger number of mutant plaques; however, we were limited by the relatively low absolute number of mutant plaques compared to the number achieved when testing a chemical mutagen. Sequencing more mutants may have allowed for a more robust analysis of the SBS signatures. This is because the C>T transition was fairly dominant, making it possible that it masked associations with other COSMIC SBS signatures. Future experiments could use reduced forms of folate, such as the naturally occurring 5-methylTHF, to test whether it is indeed the reduction of FA to DHF by DHFR that contributes to the mutagenic potential of FA in the colon, or we could use antibiotics to test the impact of the colon microbiota on FA-induced mutations in colon. Finally, we treated mutations as a proxy for cancer risk, but did not examine cancer directly. This is important as there is evidence that normal cells accumulate somatic mutations throughout life in a manner intrinsic to each tissue, without leading to cancer^62,63^.

Ultimately, we show that FA-induced mutations are tissue-specific and FA-dose dependent. The SBS signatures identified are seen in colon cancer and ALL irrespective of diet. This suggests that FA deficiency and supplementation may promote a permissive mutagenic context in colon and bone marrow, thereby facilitating endogenous mutagenic processes in these tissues. The results of our study may explain the associations between folate and these tissue-specific cancers in humans. Further studies will be needed to empirically assess the proposed mechanisms by which FA supplementation and deficiency induce tissue-specific mutations.

## Methods

### Mice

The study was approved by the Health Canada Ottawa Animal Care Committee. Mice were cared for in accordance with the Guidelines of the Canadian Council on Animal Care (CACC), described in the CACC Guide to the Care and Use of Experimental Animals^64^. A total of 60 weanling male mice derived from the Health Canada in-house MutaMouse (BALB/c x DBA/2 CD2F1) colony were used. Mice were housed at standard humidity and temperature with a 12-hour light cycle and had *ad libitum* access to food and water. At 5 weeks of age, mice were randomly assigned to one of three FA-defined diets based on the AIN-93G formula and maintained on the diet for 20 weeks (Supplemental Figure 1)^29,34^. The diets contained either 0 mg FA/kg (deficient), 2 mg FA/kg (control) or 8 mg FA/kg (supplemented) (Dyets, Inc.; Bethlehem, PA)^34^. After 10 weeks on the FA defined diets, half of the mice from each diet group (n = 10 per diet) were given a 50 mg/kg dose of ENU by gavage and the other half (n = 10 per diet) were given saline by gavage. The mice were continued on their assigned diets for another 10 weeks.

### Tissue Collection

Mice were euthanized under isoflurane anesthesia by cardiac puncture followed by cervical dislocation. Bone marrow and the colonic epithelium, referred to as “colon” hereafter, were collected. These tissues were chosen for analysis because they are associated with folate-associated pathologies, including megaloblastic anemia and colon cancer. The full-length colon was dissected, fecal pellets were gently squeezed out and it was flushed with 5 mL of cold PBS. It was cut open and laid flat with the lumen side facing up on a glass plate. The epithelial layer was lightly scraped away from the underlying colon tissue using two glass sides and collected into a 1.5 mL Eppendorf tube. Bone marrow collection was conducted as previously described^34^. Tissues were flash frozen in liquid nitrogen and stored at −80°C.

### DNA Extraction

DNA was extracted from frozen colon epithelium using the phenol/chloroform extraction procedure as previously described^65^. The steps for ‘Small intestine and colon’ extraction were followed, skipping the cracking step. Briefly, the colon was thawed and digested in 5 mL of lysis buffer and incubated at 37°C overnight, with gentle shaking. Following digestion, RNase A was added to a final concentration of 100 ug/mL and incubated for 1 h at 37°C. After addition of 5 M NaCl to a final concentration of 1.5 M, the sample was mixed and centrifuged at 2,000 x *g* for 20 min. The aqueous phase was transferred to a new 15 mL tube. Hydrated phenol was equilibrated with an equal volume of 1 M Tris-HCl, pH 8, and shaken, and the upper aqueous phase was discarded; this was repeated with 0.1 M Tris-HCl, pH 8. An equal volume of chloroform:isoamyl alcohol (24:1) was added to the phenol, mixed and centrifuged for 10 min at 1500 x *g*. An equal volume of the organic phase was added to each sample, and samples were inverted for 20-30 min and then centrifuged at 1500 x *g* for 10 min. The aqueous phase was transferred to a new 15 mL tube leaving behind any white precipitate. 5 M NaCl was added to each sample to bring the concentration to 200 mM (1/50 volume) and mixed. An equal volume of chloroform/isoamyl alcohol was added to the samples, then again, inverted for 20-30 min and then centrifuged at 1500 x *g* for 10 min. The aqueous fraction was transferred to a new 15 mL tube followed by the addition of ethanol at a 2:1 ratio to start DNA precipitation. DNA was spooled using a glass rod and washed with 70% ethanol. The sample was dried and then dissolved in TE buffer (10 mM Tris pH 7.6, 1 mM EDTA). Isolated DNA was kept at 4°C until it was used for the *lacZ* assay. DNA extraction of the bone marrow had been performed previously^34^.

### MutaMouse *lacZ* Mutant Analysis

The MutaMouse transgenic mouse model has approximately 29±4 copies in tandem repeats of the lambda bacteriophage vector (*λgt10lacZ*) in chromosome 3^66^. This allows for testing the *in vivo* mutagenicity of extrinsic factors^24^ The lacZ assay was performed following the method previously described^65^. In short, the lambda vector, was recovered and packaged in vitro into lambda phages using the Transpack Packaging extract (Agilent Technologies, Santa Clara, CA). These particles are used to infect an appropriate strain of *E. coli* (*galE-lacZ*-). Mutant identification is achieved by an overnight incubation of *E. coli* on P-Gal medium permitting the growth of a bacterial lawn allowing the visibility and quantification of plaques. P-gal is toxic to *galE*-strains that express a functional *lacZ* gene; therefore only phages that have a mutated *lacZ* gene will be able to form plaques in the P-gal selective medium^31^. Non-selective titre plates serve as a control to determine the number of background plaque forming units (PFUs). The MF was calculated per sample as the number of mutant plaques divided by the total number of PFU. A minimum of 110,000 total PFUs were scored per animal. Replicate plates with MF outside the error distribution of the diet group were considered outliers and removed from the analysis. Similar methods were used for the collection, DNA extraction and *lacZ* assay of bone marrow^34^.

### *lacZ* MF Statistical Analysis

The colon *lacZ* MF data were fit to a generalized linear model with a Binomial distribution to account for over-dispersion (variability) using the *glm* function in R of the data. Three colon DNA samples did not produce the minimum PFUs (110,000) and were removed from the analysis. In addition, three samples were identified as statistical outliers based on MF using Grubb’s test. Together, a total of six of the original 60 samples were removed from the analysis. Ultimately, among the saline treated samples, a total of 10 samples each from the deficient and control FA diet groups, and 8 samples from the supplemented FA diet were included in the analysis. Among the ENU treated samples, a total of 9, 10 and 7 in the FA deficient, control and supplemented diets, respectively, were included in the analsysi (Table 1). Differences among diets within a treatment group were identified using a one-way ANOVA and Holm-Sidak pairwise multiple comparison test. Differences between tissues and diets were identified using two-way ANOVA and Holm-Sidak pairwise multiple comparison test.

### Next Generation Sequencing (NGS)

Mutant plaques were collected using a transfer pipet into sterile microtubes containing autoclaved milliQ sterile water (0.3 mutants/μL; 1 sample per tube). The mutant plaques for each mouse were pooled and stored at −80°C until sequencing. The steps described in Beal et al., 2015^61^ under “*lacZ* Mutant Plaque Collection and Preparation for Next-Generation Sequencing” were followed to amplify the lacZ plaques and purify the PCR products with the following modifications: The final volume of PCR mastermix per well was 50 μL and contained: a 10 μL aliquot from the plaque suspension, 10 μL of 5X Q5 Reaction Buffer, 0.5 μL of Q5 High-Fidelity DNA Polymerase, 1 μL of dNTPs (10 mM) (NEB Inc.), 2.5 μL of the forward primer (10 μM), 2.5 uL of the reverse primer (10 μM) and 23.5 μL of nuclease-free water. Once the PCR products were purified using the QIAquick PCR purification kit (Qiagen), the DNA was quantified using a Qubit™ dsDNA HS Assay kit (Invitrogen™) in a Qubit 2.0 Fluorometer. Samples were diluted with nuclease-free water to a final DNA concentration of 0.2 ng/μL and stored in LoBind tubes at −20°C until library preparation.

Sequencing libraries were prepared using the Nextera XT DNA Library Prep Kit (Illumina). The Nextera XT protocol to create sequencing libraries was followed except for the step where the libraries are normalized. Briefly, the Nextera transposome is used to “tagment” the genomic DNA (gDNA) of each sample, this fragments the gDNA and tags the sequences. The “tagmented” DNA is amplified in the next step using a 12 cycle PCR program. During this amplification, the Index 1 (N7XX) and Index 2 (S5XX) and full adapter sequences are added to the “tagmented” DNA. The SPRIselect reagent beads (Beckman Coulter, Inc.) replaced the AMPure XP beads, used in the protocol, to purify the libraries and remove shorter fragments. In place of the normalization steps indicated in the Nextera protocol, we assessed the quality of the libraries using a High Sensitivity DNA Screen Tape using an Agilent 2200 TapeStation. The region was set to 150-1250 bp (base pair) and its molarity (pM/L) was calculated. The libraries were diluted to a final molarity of 500 pM. Finally, 4 μL aliquots from the 500 pM solutions of each library were pooled. NGS of the *lacZ* mutant plaques was performed in-house using the Illumina NextSeq 500 System. We used the NextSeq 500/550 High Output Kit 2.5 (150 cycles) to sequence the libraries. A total of 28 samples from colon and 29 samples from bone marrow were sequenced in duplicate. Titre plaques from control diet samples of each tissue were also sequenced as negative controls for sequencing because no mutations are expected in these plaques^61^.

### Bioinformatics

Raw sequence data was converted to FASTQ format using the bcl2fastq conversion software v2.17.1.14 (Illumina, Inc) and analysed using the pipeline described in Beal et al., 2015^61^. Reads were trimmed and aligned the reference *lacZ* sequence from the MutaMouse [GeneBank: J01636.1] using bowtie2^67^. Alignment pileup for each sample was done using SAMtools^68^. Following background correction, mutations were called if they were present in both replicates and its frequency was above the calling threshold (1/# plaques sequenced of sample). Recurrent mutations, occurrence of the same mutation at the same location more than once per animal, were considered to be the result of clonal expansion and were thus counted as one independent (unique) mutation. Mutation profiles, the distribution of mutation types in the *lacZ* gene for each FA diet, was determined using only independent mutations. Significant differences in mutation profiles were determined by one-way ANOVA followed by Tukey’ s Honest Significant Difference (Tukey HSD).

### Mutation Signature Analysis

FA-induced signatures were obtained using a R script written in house and described in Beal et al. 2019^38^. Mutations for all FA diets of both tissues were imported into R with the *lacZ* coding sequence as reference to determine the 3’ and 5’ bases directly adjacent to the base substitution in order to determine the signature of each FA diet. This trinucleotide mutation context was obtained using the “mutationContext” and “motifMatrix” functions from the “SomaticSignatures” R package^69^. To compare the *lacZ* FA-induce signatures to the COSMIC SBS signatures, derived from human mutation data, they were first normalized to *lacZ* trinucleotide frequencies. This was achieved using the ratio of trinucleotide frequencies in *lacZ* to those in the human genome^38^. The cosine similarity was used to measure the similarity/closeness between FA signatures and COSMIC SBS signatures, where a value of 1 depicts an identical signature and a value of zero a completely different signature. Analysis and visualization were conducted in R version 3.5.3^70^.

### Data availability

Sequencing reads for *lacZ* mutations will be submitted to the NCBI Sequence Read Archive.

## Author contributions

S.D.G. performed colon MF and sequencing experiments and data analysis, D.L. performed bone marrow MF experiments, R.G. aided with sequencing experiments and data analysis, N.B aided with experiments, F.M. contributed to experimental design, supervised the MutaMouse experiments and sequencing, and data interpretation. A.W. contributed to experimental design, and data interpretation. A.M conceived the experiment and directed the experimental work. All authors contributed to the writing of the manuscript.

