## Supplemental material for "Mutagenicity of folic acid deficiency and supplementation is tissue-specific and results in distinct mutation profiles"

**Supplementary Figure 1:** Study design. Beginning at 5 weeks of age, MutaMouse male mice (n=20/diet) were fed one of three folic acid (FA) diets: 0 mg FA/kg (deficient), 2 mg FA/kg (control) or 8 mg FA/kg (supplemented) and maintained on the diets for a total of 20 weeks. At the 10-week timepoint, mice were treated with saline or N-ethyl-N-nitrosurea (ENU), a known mutagen, to determine the effect of FA alone and its interaction, if any, with ENU on mutations.

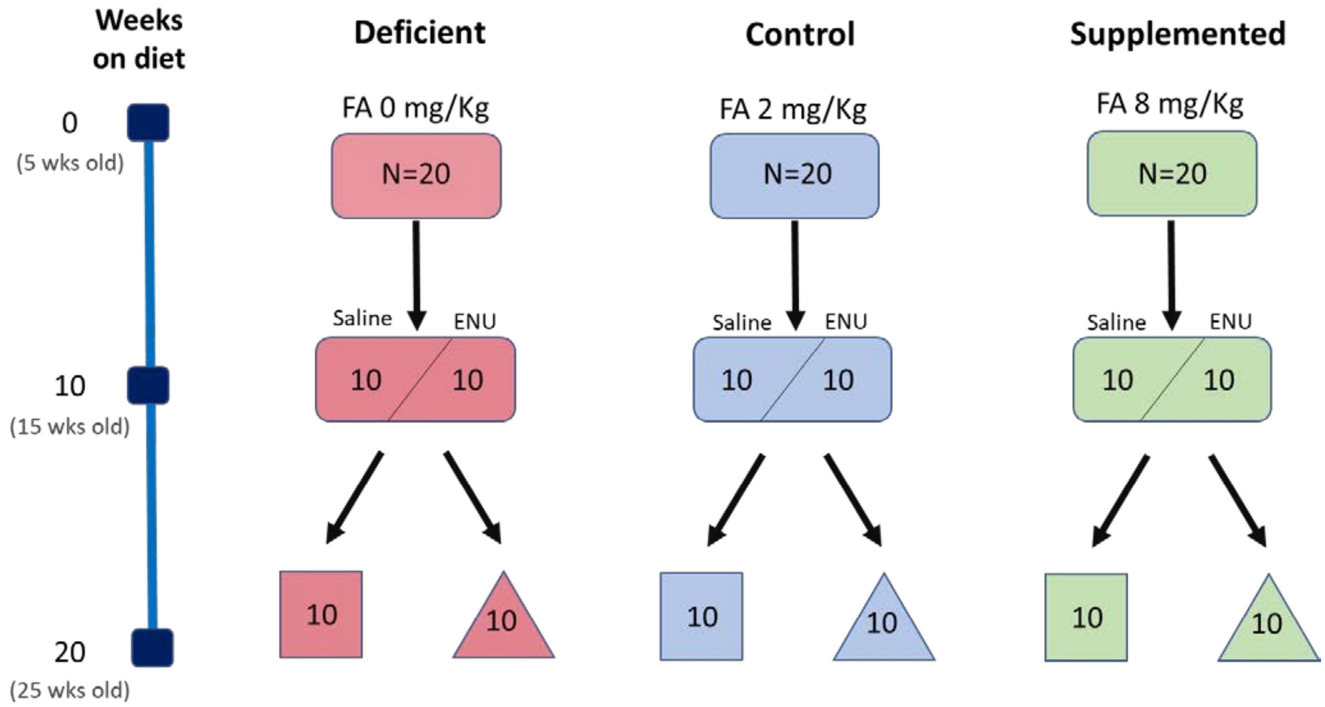

Supplementary materials:
Diaz et al. Mutagenicity of folic acid deficiency and supplementation is tissue-specific and results in distinct mutation profiles

**Supplementary Figure 2:** Cosine similarities heatmap of folic acid (FA)-induced mutational signatures with all COSMIC SBS signatures in **a)** Bone marrow and **b)** Colon.

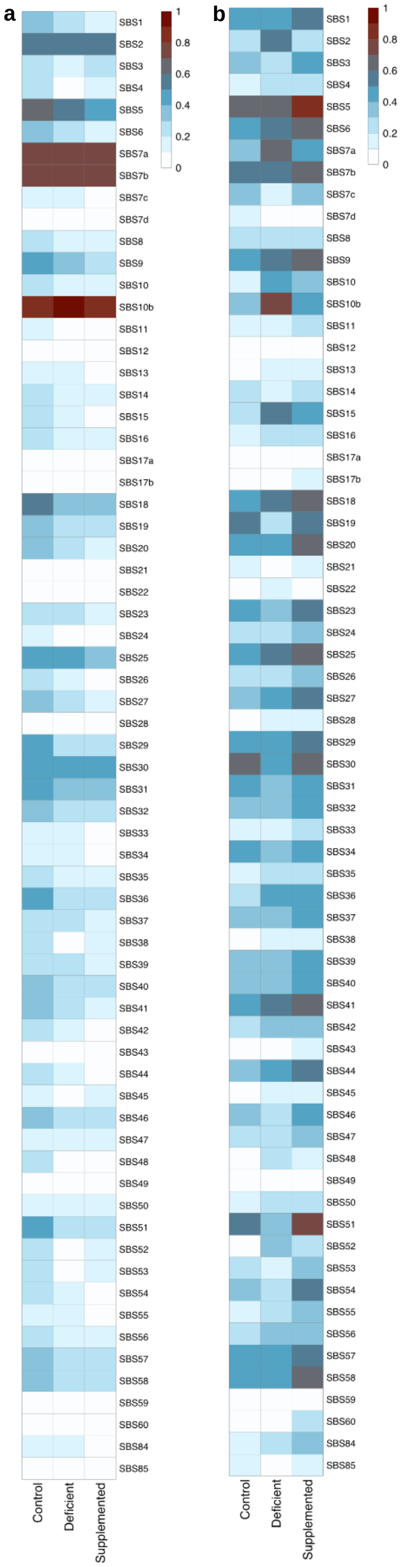

Supplementary materials:
Diaz et al. Mutagenicity of folic acid deficiency and supplementation is tissue-specific and results in distinct mutation profiles

**Supplementary Figure 3:** Bone marrow folic acid (FA) deficient signature and *lacZ* corrected COSMIC SBS signatures with cosine similarity > 0.5. SBS in red have been observed in ALL.

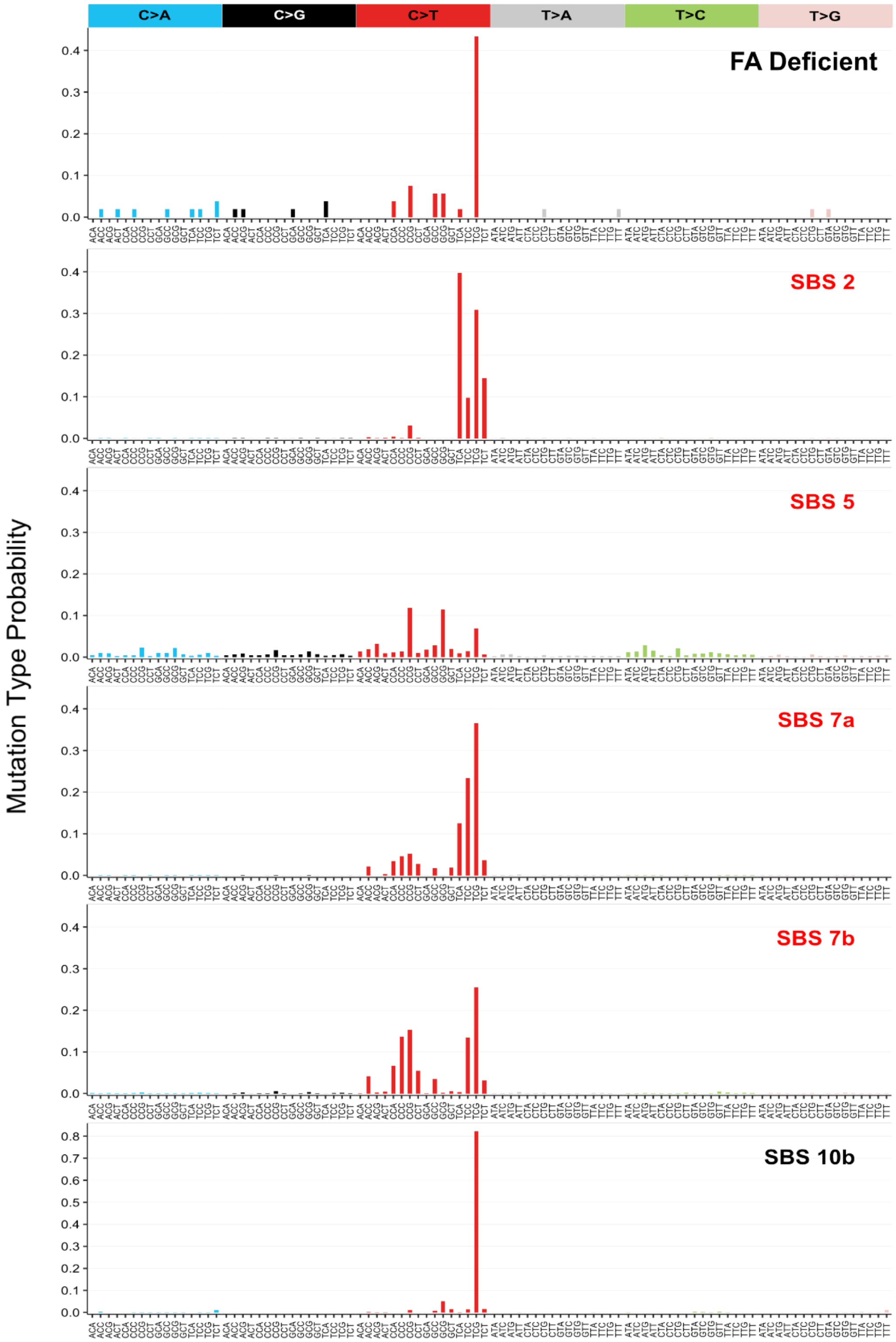

Supplementary materials:
Diaz et al. Mutagenicity of folic acid deficiency and supplementation is tissue-specific and results in distinct mutation profiles

**Supplementary Figure 4:** Colon folic acid (FA) supplemented signature and *lacZ* corrected COSMIC SBS signatures with cosine similarity > 0.5. SBS in red have been observed in CRC.

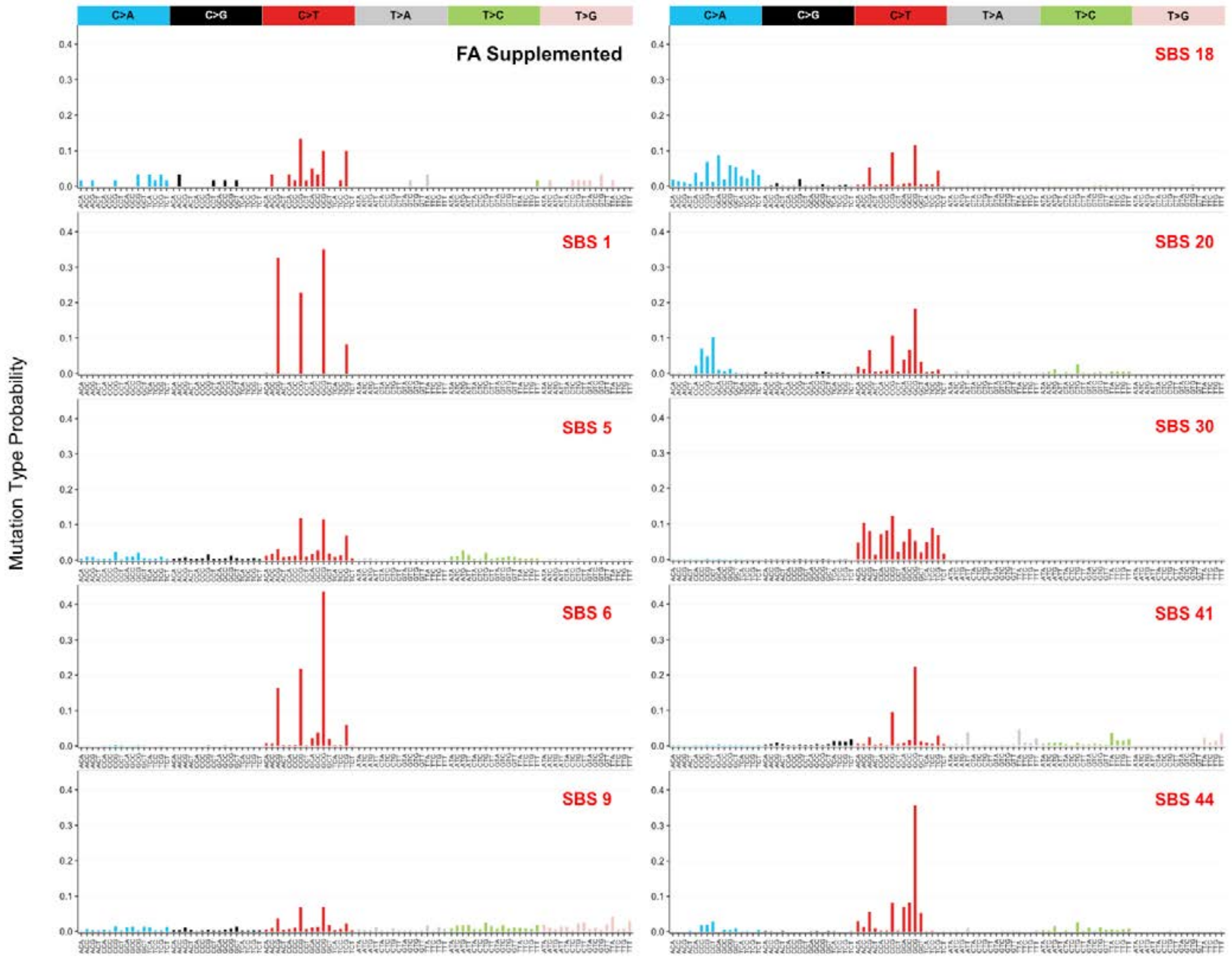

Supplementary materials:
Diaz et al. Mutagenicity of folic acid deficiency and supplementation is tissue-specific and results in distinct mutation profiles

**Supplementary Table 1: Samples sequenced, and number of independent mutations identified in the *lacZ* gene from the colon and bone marrow.** The total number of mice (samples) per folic acid (FA) diet that were sequenced, and the number of base substitutions, deletions and insertions identified per FA diet

| | | | | | | All mutations | Independent mutation types | | |
|---|---|---|---|---|---|---|---|---|---|
| **Tissue** | **FA diet** | **No. of samples** | **Total** | **Total no. of libraries (with duplicates)** | **No. of samples used in analysis[1]** | **Total** | **Base substitutions** | **Deletions** | **Insertions** |
| Bone Marrow | Deficient | 10 | 29 | 58 | 10 | 194 | 53 | 11 | 1 |
| | Control | 10 | | | 9 | 114 | 40 | 11 | 2 |
| | Supplemented | 9 | | | 8 | 92 | 28 | 3 | - |
| | | | | | **Total** | 400 | 121 | 25 | 3 |
| Colon | Deficient | 10 | 28 | 56 | 9 | 126 | 44 | 7 | 3 |
| | Control | 10 | | | 9 | 99 | 39 | 12 | 3 |
| | Supplemented | 8 | | | 8 | 136 | 52 | 5 | - |
| | | | | | **Total** | 361 | 135 | 24 | 6 |

[1]Number of samples used in the analysis may differ from the original number of samples sequenced if a sample was dropped after correcting for false mutation proportion and applying the mutation-calling threshold, as described in the methods.

Supplementary materials:
Diaz et al. Mutagenicity of folic acid deficiency and supplementation is tissue-specific and results in distinct mutation profiles

**Supplementary Table 2: Bone marrow folic acid (FA) signatures cosine similarity values with COSMIC SBS signatures.** Values in grey have a cosine < 0.5. SBS in red have been observed in acute lymphocytic leukemia.

| FA Diet | | | |
|---|---|---|---|
| **Control** | **Deficient** | **Supplemented** | **Cosmic Signatures** |
| 0.576 | 0.589 | 0.530 | SBS2 |
| 0.620 | 0.529 | 0.444 | SBS5 |
| 0.781 | 0.791 | 0.752 | SBS7a |
| 0.788 | 0.750 | 0.739 | SBS7b |
| 0.868 | 0.958 | 0.898 | SBS10b |
| 0.517 | 0.345 | 0.341 | SBS18 |

Supplementary materials:
Diaz et al. Mutagenicity of folic acid deficiency and supplementation is tissue-specific and results in distinct mutation profiles

**Supplementary Table 3: Colon folic acid (FA) signatures cosine similarity values with COSMIC SBS signatures.** Values in grey have a cosine < 0.5. SBS in red have been observed in colorectal cancer.

| FA Diet | | | |
|---|---|---|---|
| **Control** | **Deficient** | **Supplemented** | **Cosmic Signatures** |
| 0.420 | 0.443 | 0.585 | SBS1 |
| 0.245 | 0.524 | 0.293 | SBS2 |
| 0.647 | 0.651 | 0.853 | SBS5 |
| 0.459 | 0.518 | 0.665 | SBS6 |
| 0.381 | 0.618 | 0.470 | SBS7a |
| 0.584 | 0.551 | 0.652 | SBS7b |
| 0.485 | 0.508 | 0.693 | SBS9 |
| 0.351 | 0.732 | 0.462 | SBS10b |
| 0.292 | 0.500 | 0.486 | SBS15 |
| 0.499 | 0.562 | 0.681 | SBS18 |
| 0.518 | 0.269 | 0.546 | SBS19 |
| 0.458 | 0.443 | 0.621 | SBS20 |
| 0.401 | 0.339 | 0.540 | SBS23 |
| 0.494 | 0.578 | 0.647 | SBS25 |
| 0.387 | 0.458 | 0.522 | SBS27 |
| 0.407 | 0.469 | 0.558 | SBS29 |
| 0.612 | 0.415 | 0.666 | SBS30 |
| 0.495 | 0.564 | 0.681 | SBS41 |
| 0.362 | 0.493 | 0.583 | SBS44 |
| 0.529 | 0.355 | 0.728 | SBS51 |
| 0.391 | 0.290 | 0.524 | SBS54 |
| 0.411 | 0.423 | 0.533 | SBS57 |
| 0.475 | 0.450 | 0.632 | SBS58 |
